# Intraspecific genetic variation modulates immune responses to acute heat exposure in an aquatic ectotherm

**DOI:** 10.64898/2026.06.02.729540

**Authors:** Maurine Neiman, Katri Seppälä, Dunja K. Lamatsch, Otto Seppälä

## Abstract

Climate change-induced heatwaves threaten ectotherms, whose physiology is tightly coupled to ambient temperature. Vulnerability assessments often rely on data from one or a few populations, implicitly assuming uniform thermal sensitivity across species genetic diversity. Quantifying such variation is especially important for traits with wider ecological consequences; our focus here is on immune function, which shapes disease dynamics. We addressed this knowledge gap using ten clonal lineages of the New Zealand snail *Potamopyrgus antipodarum* exposed to ambient (17°C) or heatwave conditions (27°C) for 4 or 8 days. We measured two complementary innate immune traits: general phenoloxidase-like (PO-like) activity, which integrates the activity of multiple phenoloxidase enzymes, and laccase activity, which targets a specific PO enzyme subclass important in mollusc immunity. Heat exposure suppressed both traits, but patterns differed across clones. While PO-like activity declined uniformly, laccase activity showed substantial among-clone variation in heatwave responses at day 4, though these differences converged by day 8. Heat-induced immune suppression is thus trait-specific, depends on genetic background, and varies with exposure duration. Together, these results demonstrate that studies limited to a single genotype, population, or timepoint risk miscalculating species-level vulnerability.

## Introduction

Anthropogenic climate change is increasing the frequency, intensity, and duration of heatwaves worldwide, with particularly acute consequences for ectothermic organisms whose physiology is closely coupled to ambient temperature (Sales et al. 2018, Jørgensen et al. 2022, Wang et al. 2024). Unlike gradual increases in mean temperature, heatwaves impose rapid thermal challenges that can disrupt cellular homeostasis, elevate metabolic costs, and compromise organismal performance (Harding et al. 2023, Pörtner et al. 2023). Understanding how ectotherms respond to acute thermal extremes is therefore central to predicting biodiversity outcomes under continued warming.

Assessments of species vulnerability to thermal extremes typically rely on physiological measurements from one or a few populations, implicitly assuming that thermal sensitivity is uniform across species genetic diversity (Bennett et al. 2019). This assumption is increasingly difficult to justify because the intraspecific variation in physiological responses to environmental changes can be substantial, sometimes equaling interspecific variation (Des Roches et al. 2018). Overlooking this variation can yield misleading predictions of species response to environmental change (Bolnick et al. 2011, Des Roches et al. 2018). Whether and how physiological responses to heatwaves vary across species genetic diversity is therefore a question with direct implications for vulnerability assessments.

Immune function is a key component of organismal performance and is often sensitive to changes in ambient temperature (Adamo & Lovett 2011, Dittmar et al. 2014). In invertebrates, elevated temperatures alter the activity of innate immune effectors—including phenoloxidase (PO), antimicrobial peptides, and haemocyte-mediated responses—often reducing constitutive defence at temperatures above the thermal optimum (Leicht et al. 2013, Wojda 2017). The generalizability of most such studies is nevertheless limited by their focus on single populations or laboratory strains (Sandmeier 2024), despite accumulating evidence that intraspecific variation in physiological traits, including thermal responses, is widespread (reviewed in Bolnick et al. 2003, McKenzie & Anttila 2025; e.g., Nati et al. 2021, DeLiberto et al. 2022).

Direct empirical tests of how genotypic variation shapes immune responses to heatwaves remain scarce, particularly in aquatic invertebrates. The available evidence indicates that thermal effects on individual susceptibility to infections depend on genetic background in various systems (Blanford et al. 2003, Mitchell et al. 2005), and that heritable variation underlies heatwave-mediated immune changes in at least one freshwater snail species (Leicht et al. 2017). These findings suggest that genotype-by-environment (G × E) interactions are likely a general feature of immune thermal sensitivity, but their breadth, trait specificity, and temporal dynamics are poorly resolved.

Resolving these gaps requires study systems in which (i) genotypic variation can be cleanly partitioned from environmental and developmental variance, (ii) replicate individuals of the same genotype can be exposed to controlled thermal treatments, and (iii) several immune traits can be assayed in parallel. Naturally occurring obligately asexual lineages meet these criteria: each lineage represents a near-genetically uniform replicate, and lineages drawn from across a species’ range capture meaningful natural genetic diversity. The freshwater snail *Potamopyrgus antipodarum* is particularly well suited to this purpose because its natural populations harbour many independently derived asexual lineages (Dybdahl & Lively 1995, Paczesniak et al. 2013). Survivorship (e.g., Møller et al. 1994), reproductive output (Gust et al. 2011), and oxygen consumption (Greimann et al. 2020) in *P. antipodarum* are all susceptible to heat challenges, and metabolic responses to high temperatures can depend on genetic background (Matoo et al. 2023). Whether immune thermal sensitivity also varies across genetic backgrounds has not yet been tested.

We used ten clonal lineages of *P. antipodarum*, each derived from a separate natural population across the species native range in New Zealand or from an invasive North American population, to characterize how genotypic variation shapes snail immune activity under a simulated heatwave. We measured two complementary immune traits: general phenoloxidase-like (PO-like) activity, which reflects the combined oxidative potential of multiple PO enzymes, and laccase activity, a more targeted measure of one PO enzyme subclass important in mollusc immunity (Le Clec’h et al. 2016, Seppälä & Schlegel 2023). By sampling at two time points during heat exposure, we additionally assessed the temporal dynamics of these responses. We tested three predictions: (1) acute heat exposure depresses constitutive immune activity; (2) the magnitude of this thermal response varies among genotypes, indicating G × E interactions for immune thermal sensitivity; and (3) the strength of genotype-specific responses changes with exposure duration, reflecting either the accumulation of physiological constraints or genotype-specific acclimation trajectories. By testing whether and how heat-induced changes in immune activity vary across genotypes, this study addresses a fundamental gap in our ability to scale physiological measurements into species-level assessments of heatwave vulnerability and its downstream consequences for host–parasite dynamics.

## Methods

### Heat Wave Experiment

We initiated the experiment with 720 adult female *P. antipodarum* snails representing 10 distinct asexual triploid lineages. Each of these lineages was descended from a single female collected from one of 9 New Zealand lakes (Alexandrina, Te Anau, Mapourika, Okareka, Kaniere, Taupo, Heron, Gunn, Grasmere) or from an invasive population originating from Polecat Creek, Wyoming, USA (US1). These locations are known to harbour genetically distinct *P. antipodarum* populations (Paczesniak et al. 2013, Donne et al. 2020). All lineages had been maintained under common laboratory conditions for more than 10 years, minimizing the influence of environmental differences associated with collection sites. Because growth in female *P. antipodarum* stops at reproductive maturity (e.g., Winterbourn 1970), we selected the largest females from each lineage for the experiment in order to maximize the likelihood that all individuals selected had reached their full adult size.

We used a fully factorial experimental design crossing genetic lineage and temperature treatment. 36 individuals from each lineage were haphazardly assigned to each of the two temperature conditions (17 °C and 27 °C). The two temperatures were chosen to represent a baseline condition (17 °C) and an elevated temperature (27 °C) consistent with acute heatwave conditions experienced by freshwater systems within the species range (Salinger et al. 2020). For each lineage-by-temperature combination, we distributed the snails among twelve 0.23-L plastic cups filled with filtered lake water, with three individuals per cup. We housed the cups in temperature-controlled water baths corresponding to their assigned temperature treatment, and water baths were maintained in a room set to a 24-h light cycle with simulated dawn (07:00–07:30), daylight (07:30–19:30), dusk (19:30–20:00), and darkness (20:00–07:00). At the start of the experiment, each cup received a pinch of commercial freshwater fish food (Sera Micron Fry Food).

On day 4 of the experiment, we processed half of the snails (6 cups per lineage-by-temperature combination; 120 cups total) for downstream analyses with one snail per cup dedicated to each immune assay (PO-like and laccase activity, see below) and the remaining snail used for quantification of embryo production. For the snails to be used in immune assays, we first measured the snail size under a dissecting microscope by recording the shell length (apex to aperture) and then snap-froze the entire snail (including shell) in liquid nitrogen and stored it at -80°C until the immune assay. For the snail used to quantify embryo production, we again measured shell length and then opened the brood pouch to count embryos, if any. We then changed the water for the remaining 360 snails (120 cups) and supplied each cup with a pinch of fish food. On day 8 of the experiment, we performed the same sampling procedure for the 360 remaining snails. We chose the day 4 and day 8 timelines to capture both early and more prolonged responses to acute heat exposure while keeping the total experimental duration within the range of natural summer heatwaves in lakes within the species range (Salinger et al. 2020).

### Enzyme Assays

We quantified immune-related oxidative enzyme activity in experimental snails using spectrophotometric assays of phenoloxidase-like (“PO-like”) activity and laccase activity. In molluscs and other invertebrates, the prophenoloxidase (proPO) activating system is a key component of humoral innate immunity and contributes to oxidative and cytotoxic defence reactions such as melanization (Cerenius & Söderhäll 2021). Phenoloxidase activity comprises multiple enzyme families with partially overlapping substrate specificities, including catecholases, laccases, and tyrosinases (Sugumaran & Barek 2016). Assays based on different substrates therefore provide complementary information on the overall oxidative potential of the immune system. Specifically, L-DOPA is a nonspecific substrate oxidized by multiple PO enzymes and thus yields a composite measure of PO-like activity, whereas p-phenylenediamine (PPD) is primarily metabolized by laccases, a key phenoloxidase enzyme in molluscs (Le Clec’h et al. 2016, Seppälä & Schlegel 2023).

In contrast to earlier studies on other freshwater snails [*Lymnaea stagnalis* (Seppälä & Schlegel 2023), *Biomphalaria glabrata* (Le Clec’h et al. 2016)], we measured immune enzyme activity in *P. antipodarum* using whole-tissue homogenates rather than haemolymph or purified enzyme fractions. This approach was necessary because of the small body size of *P. antipodarum*. In both assays, measured activities reflect the combined oxidative potential of enzymes present in the homogenate that are capable of metabolizing the substrates, rather than the activity of a single purified enzyme. Consequently, PO and laccase activity should be interpreted as functional immune traits rather than as direct measures of specific enzyme concentrations.

Frozen snails were processed individually. Each snail was placed on ice and homogenized in 50 μl of ice-cold 10 mM cacodylate buffer (pH 8.4) using a disposable pestle. Homogenates were centrifuged at 5,000 g for 2 min at room temperature to remove debris, after which samples were immediately returned to ice. All subsequent steps were performed using chilled reagents to minimize enzymatic degradation prior to assay initiation. Because of the limited amount of tissue available per individual, we did not include technical replicates; each snail therefore represents a single biological replicate for each assay. Negative control samples were prepared using Milli-Q water in place of snail homogenate.

PO-like activity was measured using L-3,4-dihydroxyphenylalanine (L-DOPA) as a substrate. The L-DOPA solution was prepared by dissolving 40 mg of L-DOPA in 10 ml of 10 mM cacodylate buffer (pH 8.4), vortexed for 45 min, protected from light, and kept on ice until use. For each sample, 40 µl of homogenate supernatant was pipetted into a well of a flat-bottom 96-well microtiter plate, with control wells receiving 40 µl of Milli-Q water. Reactions were initiated by adding 120 µl of ice-cold L-DOPA solution to each well, followed by gentle mixing while avoiding bubble formation. Absorbance was then measured at 490 nm every 30 s for 40 min at 37 °C using a microplate reader (BioTek, Winooski, VT, USA; instrument measurement range: 0–4 with 0 being completely transparent and 4 non-transparent), with plates shaken in double-orbital mode for 1 s prior to each measurement.

Laccase activity was assessed using 1,4-phenylenediamine (PPD) as a substrate. The assay followed the same procedure as the PO-like activity assay, except that the substrate solution was prepared by dissolving 22 mg of PPD in cacodylate buffer, vortexed for 5 min, and absorbance was measured at 465 nm. In both assays, samples were processed and measured under identical conditions, and snails from different experimental treatments and genetic lineages were evenly distributed across microtiter plates to minimize potential plate-specific effects.

### Data Analyses

Seven snails (of 720; 0.97%), each from a separate cup, died during the experiment, and two additional snails were lost prior to data collection. To maximize sample sizes for immune enzyme assays, we treated these dead or lost individuals as those that would otherwise have been used for embryo quantification. Shell length data were missing for two individuals (Alexandrina 17°C treatment, day 8 data collection) included in the immune assays.

All statistical analyses were conducted in IBM SPSS Statistics, version 31, and figures were created using R 4.5.3 (2026-03-11) software (R Core Team 2026) and ggplot2 (Figs. 1, 2) or IBM SPSS v. 31 (Supplemental Figs. 1-4). For each snail and assay, we extracted the reaction rate (“Slope”), calculated as the slope between the starting absorbance and the earliest occurrence of the maximum absorbance value. In each assay, background absorbance was accounted for by calculating the mean of the eight control wells run on each plate and subtracting this value from the corresponding snail measurements.

**Figure 1.**
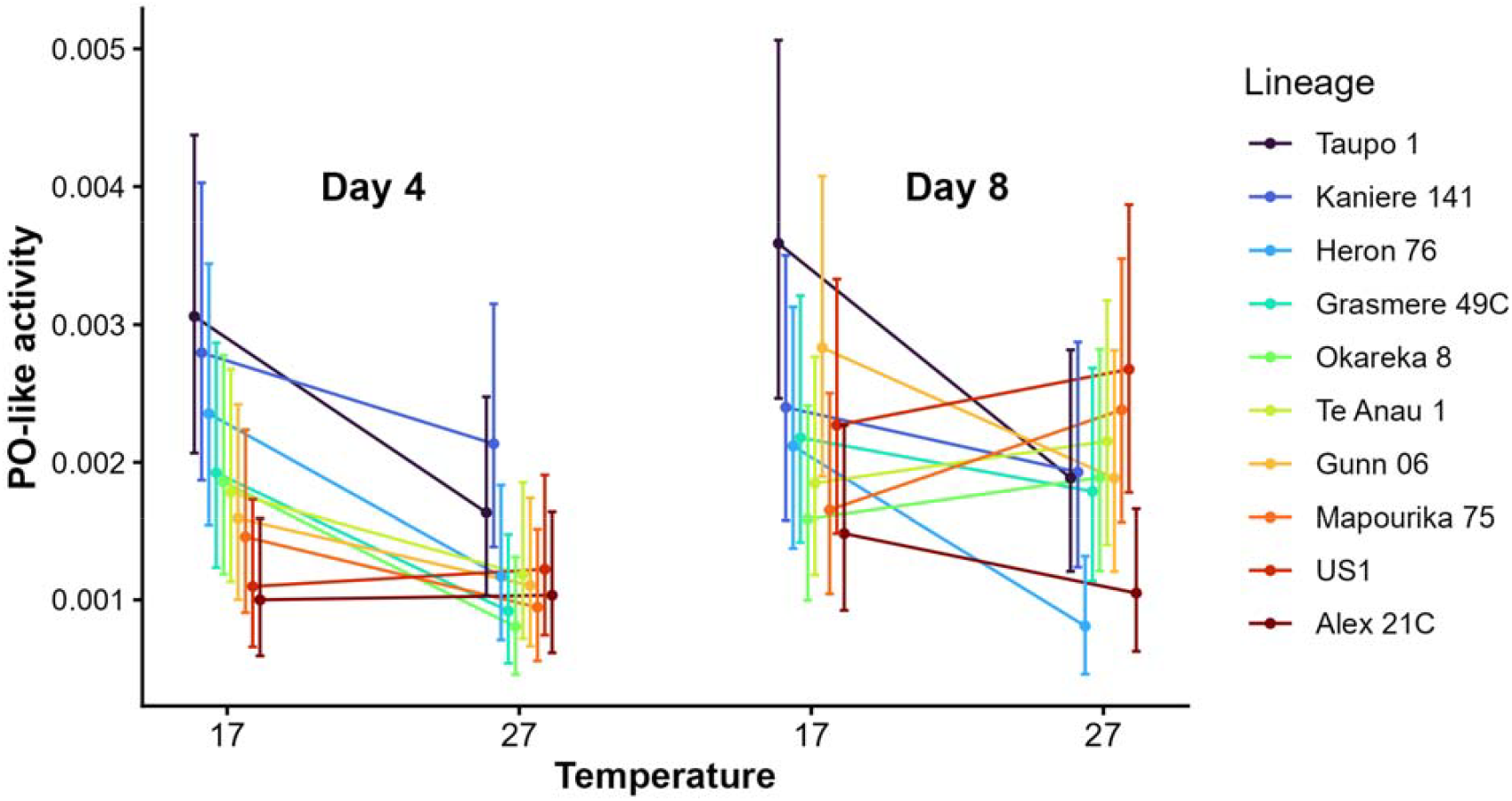
Phenoloxidase-like (PO-like) activity (reaction slope; estimated marginal mean ± 95% CI) and in 10 lineages of *Potamopyrgus antipodarum* snails maintained in different temperature treatments (17°C, 27°C) for 4 and 8 days. Lineages are arranged within each treatment combination according to their rank order at 17°C at day 4 of the experiment, and they are connected across the treatments using reaction norms at each day.

**Figure 2.**
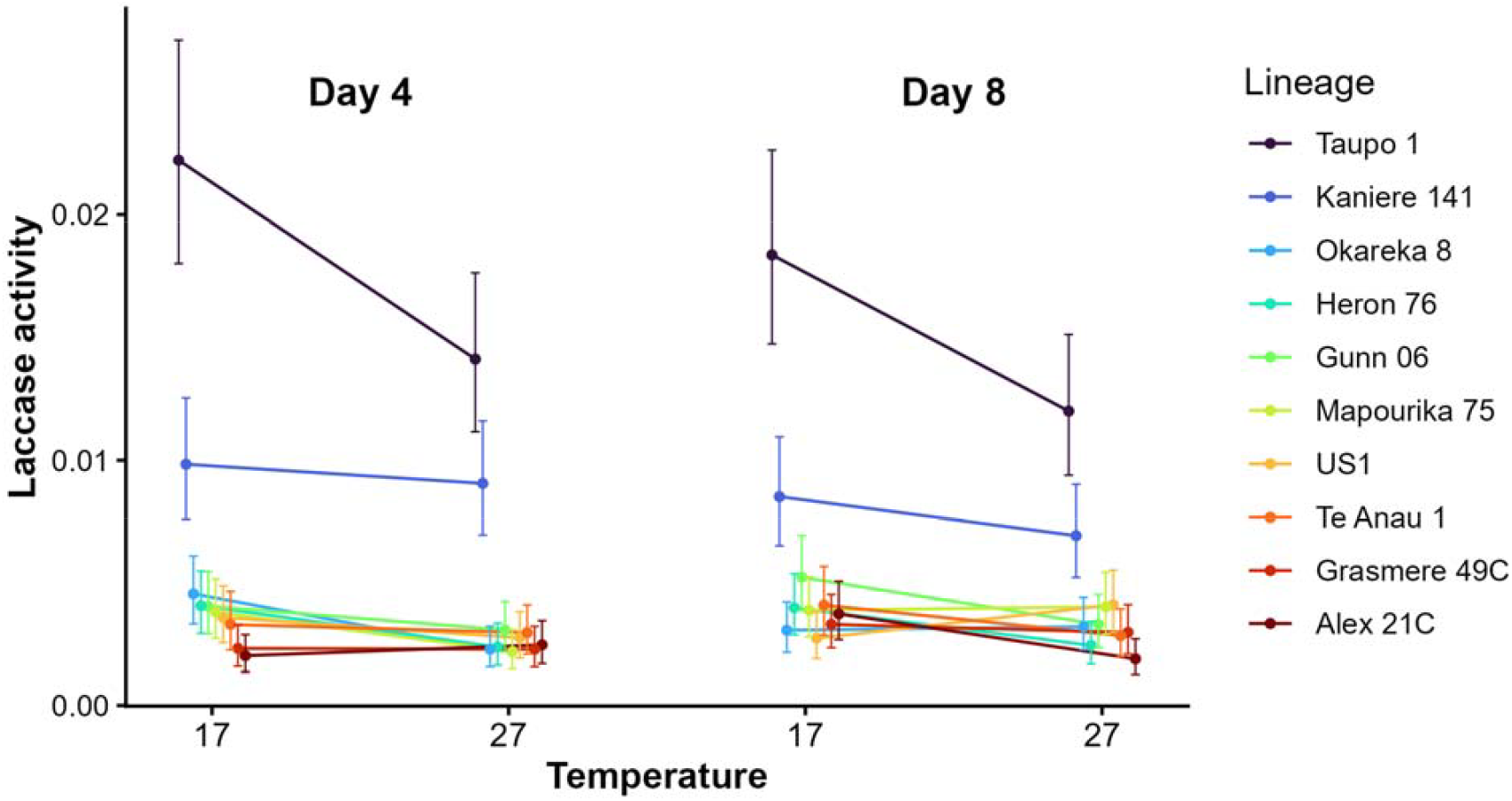
Laccase activity (reaction slope; estimated marginal mean ± 95% CI) in 10 lineages of *Potamopyrgus antipodarum* snails maintained in different temperature treatments (17°C, 27°C) for 4 and 8 days. Lineages are arranged within each treatment combination according to their rank order at 17°C at day 4 of the experiment, and they are connected across the treatments using reaction norms at each day.

We analysed the variation in PO-like and laccase enzyme activities using analyses of variance (ANOVAs) after transforming enzyme activity measures (reaction slopes) using a fourth-root transformation to homogenize error variances. For each enzyme, we initially fitted a model including temperature treatment (17°C vs 27°C) and exposure duration (4 vs 8 days) as fixed factors, and lineage as a random factor (main effects and all interactions). Additionally, assay plate (main effect only) was included as a random factor. In these analyses, we evaluated potential outliers using studentized residuals (> 4) and Cook’s and Leverage distances (> 0.5) and excluded such observations (PO-like: 0 observations; laccase: 2 observations). For laccase activity, the full model revealed a significant temperature × exposure duration × lineage interaction (see Results section). To clarify the source of this interaction, we conducted additional analyses separately for each exposure duration (day 4 and day 8). For these analyses, we fitted models including temperature, lineage, assay plate, and the temperature × lineage interaction.

Because shell length differed among lineages and could potentially contribute to lineage-specific immune responses observed for laccase (see Results), we conducted additional exploratory analyses to evaluate whether lineage effects in laccase activity were associated with variation in snail size. However, shell length and its interactions with other factors could not be included in the ANOVAs due to insufficient degrees of freedom resulting from the large number of parameters relative to the available sample size. We thus used separate Spearman’s correlations to evaluate the relationship between among-lineage variation in mean shell length and mean laccase activity. We conducted these analyses using estimated marginal means from the above ANOVA models. Only 10 snails (out of 419 dissected) produced any embryos, preventing us from analyzing this part of the dataset. Six of these snails were from the Taupo 1 lineage, 3 were from the Te Anau 1 lineage, and 1 was from the Kaniere 141 lineage, The number of embryos produced ranged from 1-14 (mean = 5.1, standard deviation = 4.358). This low frequency of embryo production is not unprecedented in these snails, which often feature highly seasonal reproduction (Winterbourn 1970).

## Results

PO-like activity was decreased by 7.3% under the high-temperature treatment (Fig. 1; Table 1) and was 7.3% higher at day 8 of the experiment compared to day 4 (Fig. 1; Table 1). Lineage or its interactions with other factors did not influence PO-like activity (Fig. 1; Table 1). Laccase activity showed a significant temperature × day × lineage interaction (Fig. 2; Table 1). This result indicates that genotypes differed in their thermal responses, an effect which was further modified by exposure duration. Separate analyses by sampling day showed that on day 4, laccase activity of individuals at the high temperature was 6.9% lower compared with those at the benign temperature (Fig. 2; Table 2). Furthermore, lineages differed in their laccase activity (Fig. 2; Table 2) and the negative effects of the high temperature (Fig. 2; Table 2). On day 8, laccase activity was 5.6% lower at the high temperature (Fig. 2; Table 2) and lineages differed from each other (Fig. 2; Table 2), but no longer as a function of temperature (Fig. 2; Table 2).

**Table 1.** Mixed-model analyses of variance (ANOVAs) on phenoloxidase-like PO-like activity and laccase activity in *Potamopyrgus antipodarum* by temperature treatment (17°C, 27°C), exposure duration (4 or 8 days after the temperature manipulation started), snail lineage (10 clones), and assay plate.

|  | Source | df | MS | F | p |
| --- | --- | --- | --- | --- | --- |
| PO-like activity | Temperature (T) | 1 | 0.140 | 10.706<sup>a</sup> | 0.010 |
|  | Day (D) | 1 | 0.012 | 8.878<sup>a</sup> | 0.015 |
|  | Lineage (L) | 9 | 0.003 | 1.525<sup>b</sup> | 0.279 |
|  | Assay plate | 5 | 0.001 | 0.741 | 0.593 |
|  | T × D | 1 | 0.002 | 2.523<sup>c</sup> | 0.147 |
|  | T × L | 9 | 0.001 | 1.775<sup>c</sup> | 0.203 |
|  | D × L | 9 | 0.001 | 1.866<sup>c</sup> | 0.183 |
|  | T × D × L | 9 | 0.001 | 1.004 | 0.438 |
|  | Error | 195 | 0.001 |  |  |
| Laccase activity | Temperature (T) | 1 | 0.016 | 17.251<sup>a</sup> | 0.002 |
|  | Day (D) | 1 | 0.001 | 0.721<sup>a</sup> | 0.418 |
|  | Lineage (L) | 9 | 0.043 | 77.922<sup>b</sup> | 0.123 |
|  | Assay plate | 5 | 0.001 | 2.430 | 0.037 |
|  | T × D | 1 | 0.000 | 0.132<sup>c</sup> | 0.725 |
|  | T × L | 9 | 0.001 | 0.744<sup>c</sup> | 0.667 |
|  | D × L | 9 | 0.001 | 0.691<sup>c</sup> | 0.705 |
|  | T × D × L | 9 | 0.001 | 2.111 | 0.030 |
|  | Error | 193 | 0.001 |  |  |

<sup>a</sup> (T × L) as the error term
<sup>b</sup> (T × L) + (D × L) + (T × D × L) as the error term
<sup>c</sup> (T × D × L) as the error term

**Table 2.** Mixed-model analyses of variance (ANOVA) on laccase activity in *Potamopyrgus antipodarum* for each sampling day (day 4 and day 8 after the temperature manipulation started) by temperature treatment (17°C, 27°C), snail lineage (10 clones), and assay plate.

|  | Source | df | MS | *F* | *p* |
| --- | --- | --- | --- | --- | --- |
| Day 4 | Temperature (T) | 1 | 0.010 | 10.170<sup>a</sup> | 0.011 |
|  | Lineage (L) | 9 | 0.027 | 27.856<sup>a</sup> | < 0.001 |
|  | Assay plate | 5 | 0.002 | 4.128 | 0.002 |
|  | T × L | 9 | 0.001 | 2.271 | 0.024 |
|  | Error | 94 | 0.000 |  |  |
| Day 8 | Temperature (T) | 1 | 0.007 | 5.238<sup>a</sup> | 0.048 |
|  | Lineage (L) | 9 | 0.017 | 13.604<sup>a</sup> | < 0.001 |
|  | Assay plate | 5 | 0.000 | 0.274 | 0.927 |
|  | T × L | 9 | 0.001 | 1.602 | 0.126 |
|  | Error | 94 | 0.001 |  |  |

<sup>a</sup> (T × L) as the error term

Spearman’s rank correlations to evaluate whether among-lineage variation in body size contributes to lineage-level variation in laccase activity (using estimated marginal means from the ANOVA models above for laccase activity, and from analogous models for shell length; analyses restricted to treatment combinations in which a significant lineage effect on laccase activity was detected) revealed positive correlations (day 4 across both temperatures: Spearman’s correlation = 0.590, *p* = 0.006; 17 °C, day 4: Spearman’s correlation = 0.681, *p* = 0.03; 27 °C, day 4: Spearman’s correlation = 0.596, *p* = 0.069; day 8 across both temperatures: Spearman’s correlation = 0.706, *p* = 0.023; Supplemental Figs. 1-4). Altogether, body size accounted for roughly 50% of the among-lineage variation in laccase activity, suggesting that lineage differences in body size contribute to, but do not fully account for, the significant lineage-level variation in laccase activity.

## Discussion

We found that a simulated heatwave altered immune function in *Potamopyrgus antipodarum* in a way that depended on both the measured immune trait and the snail’s genetic background. Phenoloxidase-like activity declined uniformly under heat exposure, with no detectable variation among genotypes. Laccase activity, by contrast, exhibited a temperature × time × lineage interaction, indicating that genotypes differed in both the magnitude and the temporal trajectory of their thermal responses. These results support the general expectation that immune responses to acute heat exposure are not uniform across the genetic diversity of a species, but they additionally reveal that the strength of this G × E pattern is trait-specific and dependent on the duration of heat exposure.

### Heatwave-imposed suppression of immune activity

Both immune traits we measured declined at the elevated temperature, consistent with a broad pattern across ectotherms in which acute heat exposure suppresses constitutive innate immunity (Adamo et al. 2012, Dittmar et al. 2014, Sandmeier 2024). Several non-mutually exclusive mechanisms could underlie this pattern. Thermal challenge elevates baseline metabolic demand and may reallocate resources away from costly immune traits toward the maintenance of homeostasis (Pörtner 2010). In particular, tradeoffs related to the physiological response to high temperatures might impact oxidative defenses (Palmer 2018). Heat exposure can also directly destabilize enzyme structure, particularly for catalytically active proteins operating near their thermal optima (Daniel et al. 2008). Finally, heat-induced changes in upstream regulatory pathways, including the prophenoloxidase activating cascade, may dampen functional output without changes in enzyme abundance per se (Wojda 2017). Disentangling these mechanisms was beyond the scope of the present work, but our findings establish that the PO-like suppression is robust across 10 genetic backgrounds we tested and is therefore unlikely to reflect responses of any single genotype.

The magnitude of the immune decrease we observed (∼7% for both traits at day 4) is moderate compared with thermal effects reported in some ectotherms (Roth et al. 2010, Karl et al. 2011). This may reflect species-specific thermal tolerance, the relatively short heatwave duration, or compensatory mechanisms not captured by our two assays. Comparable studies on freshwater snails have reported a range of effect sizes, often modulated by resource availability or co-occurring stressors (Salo et al. 2017, Seppälä et al. 2024). Whether the moderate suppression we observed translates into measurable changes in disease susceptibility under field conditions is an open question that warrants direct testing through infection experiments.

### Trait-specific responses to heat exposure

A central finding of our study is that the two PO-mediated immune traits we measured responded differently to the heatwave, pointing to differences in their regulation. PO-like activity—a composite measure capturing the activity of multiple PO enzyme families—exhibited a consistent decline at elevated temperature across genotypes, suggesting that this aggregate immune measure is conserved or canalized in its thermal response. By contrast, laccase activity, which targets a more specific subset of PO enzymes, depended on lineage, temperature, and exposure duration, pointing to context-dependent expression. This contrast aligns with the established recognition that different PO enzyme families differ in substrate specificity, regulation, and physiological role (Cerenius et al. 2008, Le Clec’h et al. 2016, Seppälä & Schlegel 2023). Composite assays may therefore mask trait-specific G × E patterns that emerge only when individual enzyme classes are measured separately.

The trait-specific G × E has biological significance. Laccases in molluscs are increasingly recognized as components of humoral immunity with roles in melanization and anti-parasite defence (Luna-Acosta et al. 2017, Le Clec’h et al. 2016). Genotypic variation in laccase thermal sensitivity—but not in aggregate PO-like activity—suggests that thermal responses of more functionally specific immune effectors may be more evolutionarily labile than those of broadly conserved oxidative pathways. We note, however, that the contrast between PO-like and laccase responses we observed here differs from the situation reported in another freshwater snail species, where genetic variation in thermal sensitivity of PO-like activity has been detected (Leicht et al. 2017). This contrast suggests that the trait specificity of immune G × E may itself vary among species and underscores the value of measuring multiple immune traits in parallel.

### Genetic variation in immune activity

Genotypes differed substantially in laccase activity, both at baseline and in their response to heat. This finding adds to a growing body of work demonstrating genetic variation in invertebrate immune function (e.g., Cotter et al. 2004, Leicht et al. 2017, Spaan et al. 2022), and, to our knowledge, represents the first such report for *P. antipodarum*. Documenting genotypic variation in this species is of particular interest because *P. antipodarum* is a well-established model for host-parasite coevolution (e.g., Dybdahl & Lively 1998, Koskella & Lively 2007, Gibson et al. 2016) and because the characterization of G × E interactions in thermal plasticity is essential for predicting evolutionary potential under climate change (Chevin et al. 2013, Merilä & Hendry 2014).

Our experimental design, featuring one lineage per each of 10 distinct lake populations, thus cannot directly assess whether the genotypic variation we observed reflects standing genetic variation within any single snail population. Even so, among-lineage differences capture biologically meaningful heterogeneity in physiological performance across heterogeneous environments (Bolnick et al. 2011). Such heterogeneity matters for two reasons. First, it indicates that single-genotype or single-population studies risk over-or underestimating the true range of immune responses present across a species’ genetic diversity, an issue with direct relevance to climate-vulnerability assessments that scale up from physiological measurements to species-level projections (Bennett et al. 2019). Second, where comparable variation also exists within populations, it provides the raw material on which selection can act under environmental change (Orr & Unckless 2014, Uecker et al. 2014).

The temporal pattern we observed adds an additional layer of insight. Lineage × temperature interaction was significant on day 4 but no longer significant on day 8, even though the main effects of lineage and temperature remained significant. In other words, lineages differed in their thermal responses early in the heatwave, but those differences converged when the exposure continued. This may be either because longer exposure accumulates physiological constraints that override initial genetic differences (Dhabhar 2018, Alotiby 2024) or because genotypes initially employing different strategies converge on a common stress phenotype. Thus, fast and slow responders may eventually reach a similar endpoint. Distinguishing among these possibilities will require finer-grained temporal sampling (see Leicht et al. 2013). However, our findings demonstrate that snapshot measurements at a single time point can substantially mischaracterize the genotypic structure of responses to environmental change.

### Body size and immune function

Exploratory analysis showed moderate positive correlations between lineage mean body size and laccase activity, with body size accounting for roughly half of the among-lineage variation. Previous studies have demonstrated genetic variation for adult body size in *P. antipodarum* (Larkin et al. 2016, Donne et al. 2022), so these correlations suggest that heritable differences in body size, or in traits correlated with body size, contribute to, but do not fully account for, among-lineage variation in laccase activity. A direct role of body size for immune function would be more clearly indicated if immune activity also correlated positively with size *within* clonal lineages. We did not have sufficient statistical power to test this possibility, and such a test would comprise a useful next empirical step.

The pattern that larger lineages tended to show stronger immune activity is consistent with studies from other invertebrates demonstrating a positive intraspecific relationship between body size and immune function (e.g., Otterstater & Thomsen 2006, Vogelweith et al. 2013, Körner et al. 2017). The only similar example we are aware of in snails comes from *Cornu aspersum*, where the relationship between body size and immune response depended on the specific immune mechanism considered (Russo & Madec 2011).

### Implications for forecasting climate vulnerability

Our results have direct implications for how immune thermal sensitivity is incorporated into climate-vulnerability assessments. Most current frameworks rely on physiological measurements from one or a few genotypes or populations and assume that the measured response represents the species (Bennett et al. 2019). Our findings suggest that for some traits, this assumption may be justified (aggregate PO-like activity declined consistently across genotypes), but for others, it can be misleading. Where G × E thermal sensitivity is present, single-genotype estimates may either underestimate or overestimate species-level vulnerability, depending on which genotype was investigated. Beyond mean responses, the variance among genotypes is itself biologically informative: a species with substantial standing variation in immune thermal sensitivity has a different evolutionary outlook than one with little variation, even if their mean responses are identical (Chevin et al. 2013, Merilä & Hendry 2014). Our results therefore argue for greater integration of intraspecific variation—including measurement of multiple genetic backgrounds and reporting of among-genotype variance components—in eco-immunological studies of climate change.

These implications may extend to host-parasite dynamics. Heatwave-imposed immune suppression has been linked to increased susceptibility to infections (e.g., Palmer et al. 2010) and even altered host-parasite coevolutionary dynamics in other systems (Hector et al. 2023). *P. antipodarum* is well known for its coevolutionary relationship with the trematode *Atriophallophorus winterbourni*, with parasites showing local adaptation to specific snail genotypes (e.g., Lively & Dybdahl 2000, King et al. 2009). The genotype-specific immune responses we observed raise the possibility that heatwaves could alter the outcomes of these coevolutionary interactions, with downstream consequences for the maintenance of sexual vs. asexual reproduction in *P. antipodarum* (Lively 1987, Jokela et al. 2009). Direct tests of this possibility would be a valuable next step.

### Caveats & Limitations

Several caveats should be noted. First, we measured immune activity in whole-body homogenates rather than haemolymph. While this approach was necessary given the small body size of *P. antipodarum*, it means our measurements integrate enzyme activity across multiple tissues, some of which may not contribute to immune defence in the same way as circulating haemolymph. Whether the genotype × temperature interaction we observed for laccase activity is driven by tissue-specific or systemic responses therefore cannot be resolved from our data. Second, we did not measure infection outcomes directly; whether the immune changes we documented translate into altered disease susceptibility remains to be tested experimentally. Third, our two-timepoint design captures short-term dynamics but cannot resolve longer-term acclimation or recovery trajectories. Fourth, we were unable to perform planned comparisons assessing evidence for heat stress via impacts on reproduction because only a handful of snails produced any embryos. Finally, laboratory heatwaves cannot fully replicate the complexity of natural heatwave events, which may co-occur with other abiotic and biotic environmental changes such as hypoxia and resource limitation (Verberk et al. 2016, Seppälä et al. 2024). Therefore, our results should be interpreted as a controlled test of single-stressor effects rather than a forecast of field outcomes.

## Conclusions & Next Steps

We found that acute heat exposure suppresses immune activity in *P. antipodarum* in an enzyme and genotype-dependent manner. Aggregate phenoloxidase-like activity declined uniformly across genotypes, while laccase activity exhibited substantial G × E interaction that was further modulated by exposure duration. These findings indicate that *P. antipodarum* harbours among-genotype variation in thermal responses for at least some components of immune function. Although our experimental design cannot distinguish standing within-population genetic variation from among-population differences, the observed heterogeneity captures biologically meaningful physiological variation across environments (Bolnick et al. 2011), with direct relevance to ecological resilience and tolerance.

Important next steps include linking immune activity to direct measurements of infection outcomes under heatwaves (Hector et al. 2023), characterizing within-population genetic variation to assess evolutionary potential to a warming climate, and identifying the molecular pathways underlying genotype-specific differences in thermal sensitivity. More broadly, our findings highlight the importance of incorporating intraspecific genetic variation and temporal dynamics into eco-immunological studies to accurately forecast how ectotherms will cope with a warming world.

## Acknowledgements & Funding

We would like to thank Winnie Gavin for snail maintenance and Tia Philibert for help with data visualization. We are grateful for funding from the University of Innsbruck to DKL and a UIBK Guest Professorship to MN, the University of Iowa International Programs, and the Austrian Science Fund (grant no. P 34687).

## Notes

### Competing Interest Statement

The authors have declared no competing interest.

