## Supplemental Tables 1-4 for "Intraspecific genetic variation modulates immune responses to acute heat exposure in an aquatic ectotherm"

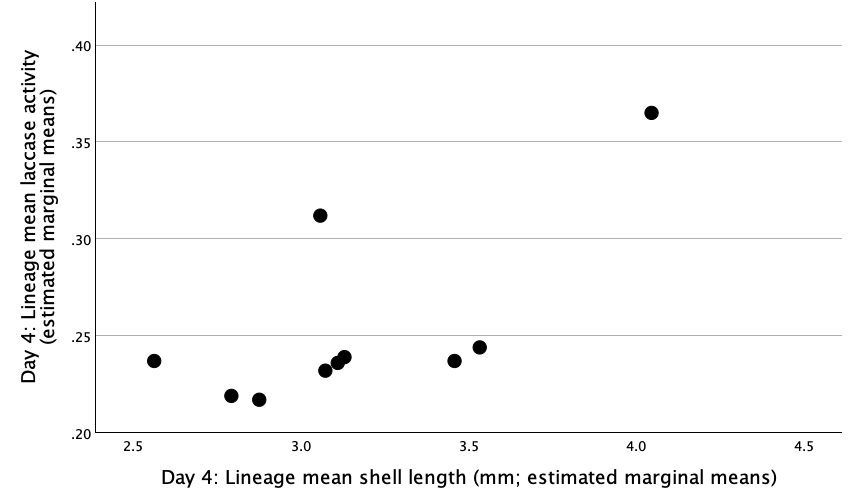

Supplemental Figure 1. There was a positive relationship between lineage mean shell length and mean lineage laccase activity on day 4 of the experiment across both temperatures (Spearman’s correlation = 0.590, *p* = 0.006). Estimated marginal means were used for this analysis and figure.

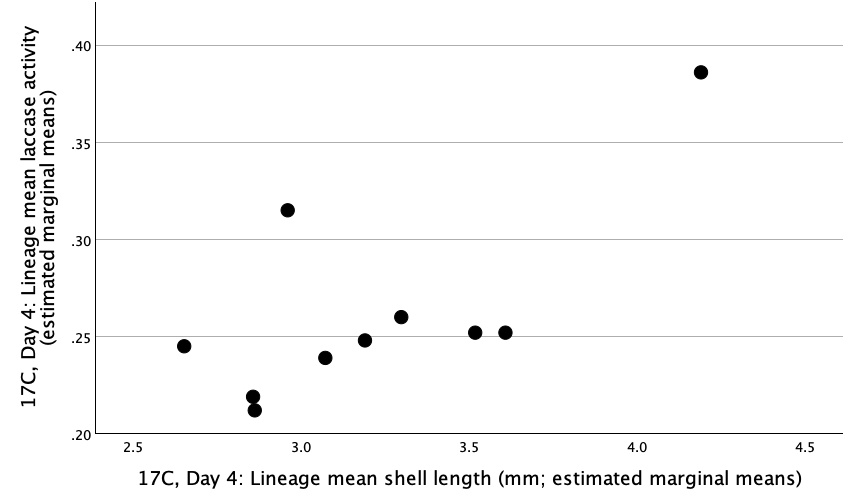

Supplemental Figure 2. There was a positive relationship between lineage mean shell length and mean lineage laccase activity on day 4 of the experiment at 17C (Spearman’s correlation = 0.681, *p* = 0.03). Estimated marginal means were used for this analysis and figure.

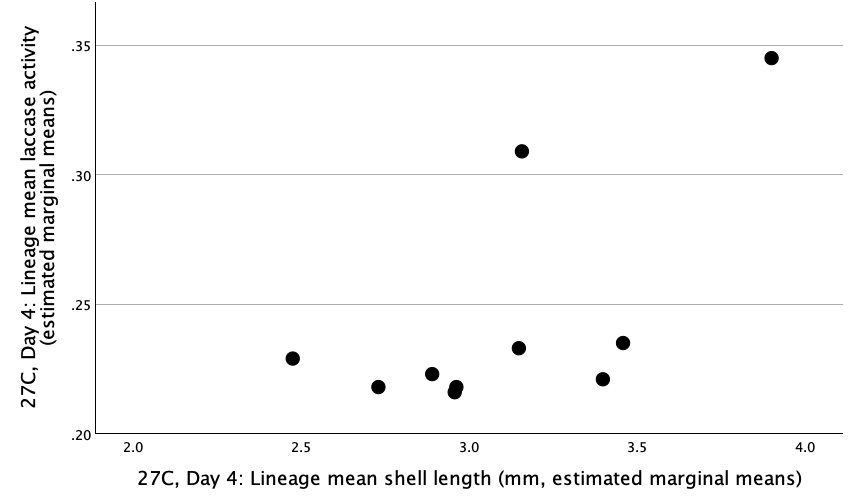

Supplemental Figure 3. There was a positive relationship between lineage mean shell length and mean lineage laccase activity on day 4 of the experiment at 27C (Spearman’s correlation = 0.596, *p* = 0.069). Estimated marginal means were used for this analysis and figure.

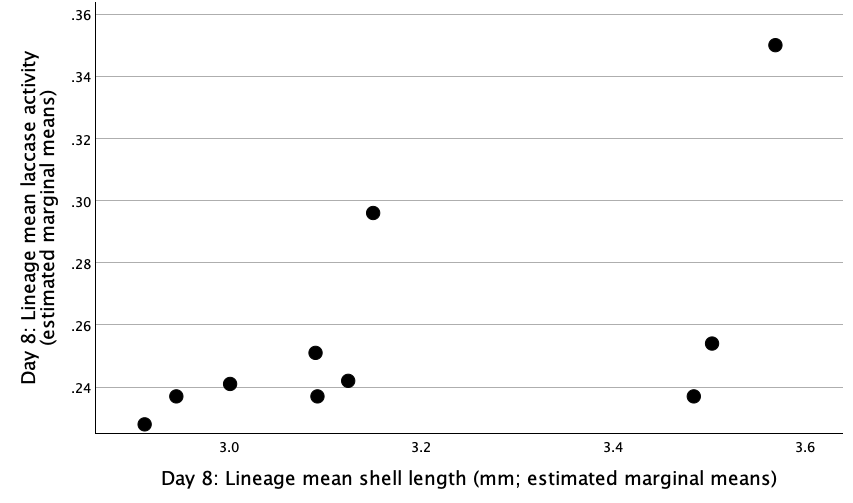

Supplemental Figure 4. There was a positive relationship between lineage mean shell length and mean lineage laccase activity on day 8 of the experiment across both temperatures (Spearman’s correlation = 0.706, *p* = 0.023). Estimated marginal means were used for this analysis and figure.
